## Supplemental Figures and legends for "Sperm-specific COX6B2 enhances oxidative phosphorylation, proliferation, and survival in lung adenocarcinoma"

A.

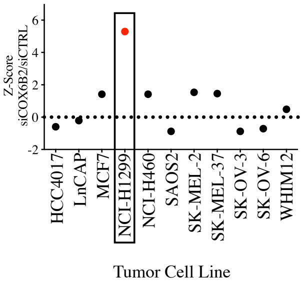


B.

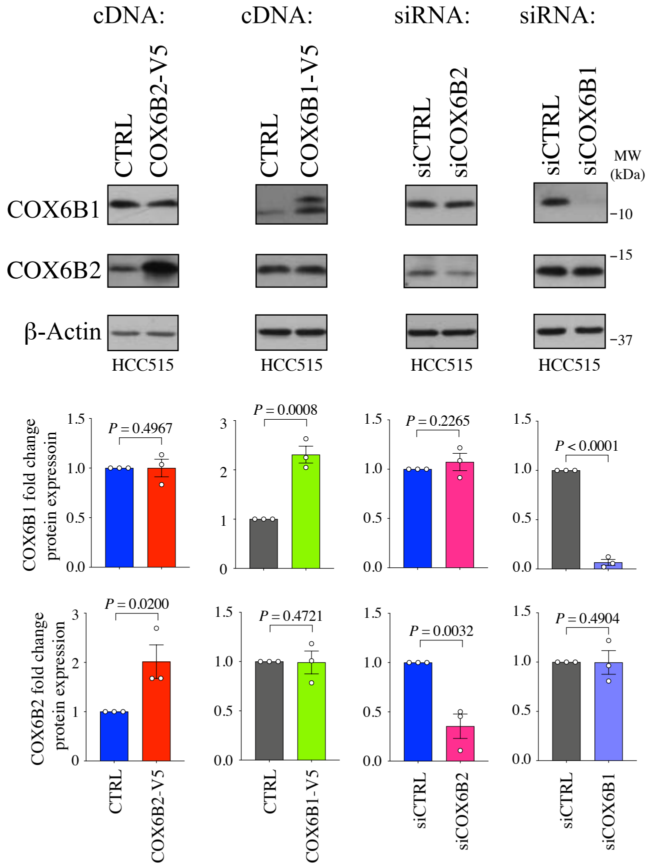


C.

| No | Pathology | Stage |
|---|---|---|
| 1 | Adenocarcinoma: 40% acinar, 30% papillary, 20% lepidic, 10% micropapillary | IB |
| 2 | Adenocarcinoma 90% solid with 10% acinar subtype | IIB |
| 3 | Solid adenocarcinoma with mucin (AB-PAS only) Nearly 100% solid | IIA |
| 4 | Adenocarcinoma, solid 60%, acinar 40% | IIIB |
| 5 | Adenocarcinoma, papillary-predominant (papillary - 85%; micropapillary - 15%) | IIB |
| 6 | Adenocarcinoma, acinar/cribriform (60%), solid (40%) | IIIA |
| 7 | Adenocarcinoma, acinar predominant | IB |
| 8 | Adenocarcinoma | IIB |
| 9 | Adenocarcinoma, solid subtype | IIA |
| 10 | Adenocarcinoma | IIB |

D.

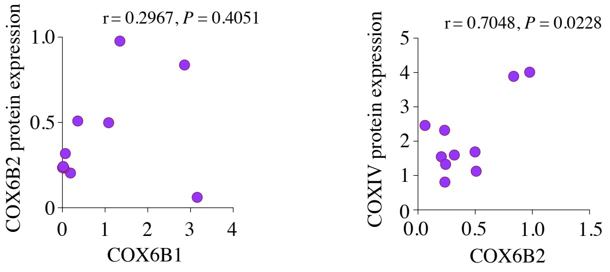

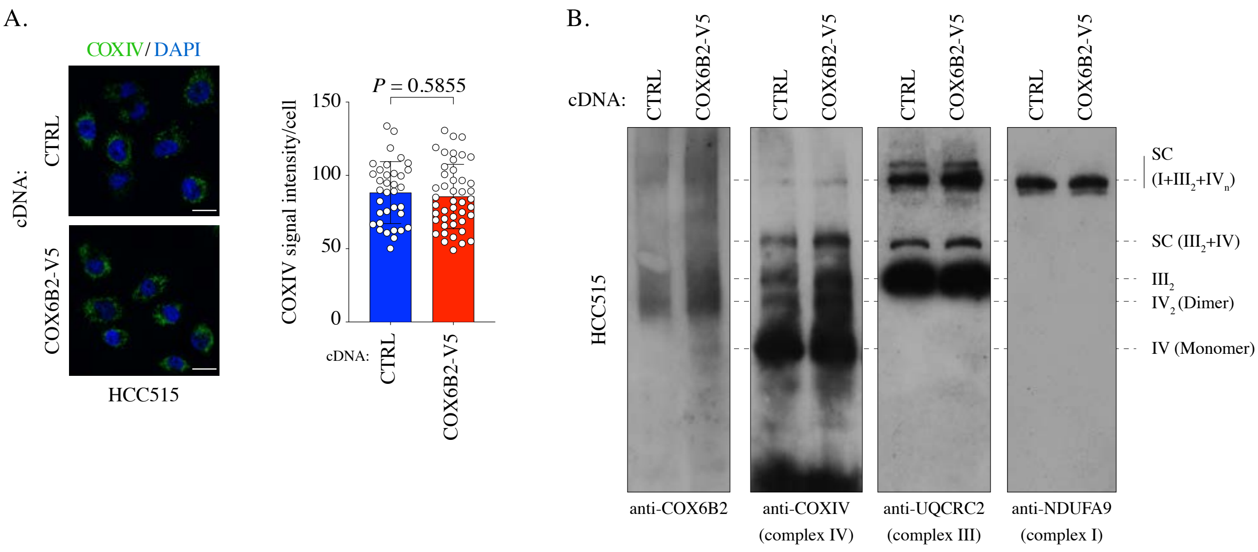
A.
COXIV / DAPI
cDNA:
CTRL
COX6B2-V5
HCC515
P = 0.5855
COXIV signal intensity/cell
150
100
50
0
cDNA:
CTRL
COX6B2-V5
B.
cDNA:
CTRL
COX6B2-V5
HCC515
SC
(I+III2+IVn)
SC (III2+IV)
III2
IV2 (Dimer)
IV (Monomer)
anti-COX6B2
anti-COXIV
(complex IV)
anti-UQCRC2
(complex III)
anti-NDUFA9
(complex I)

!

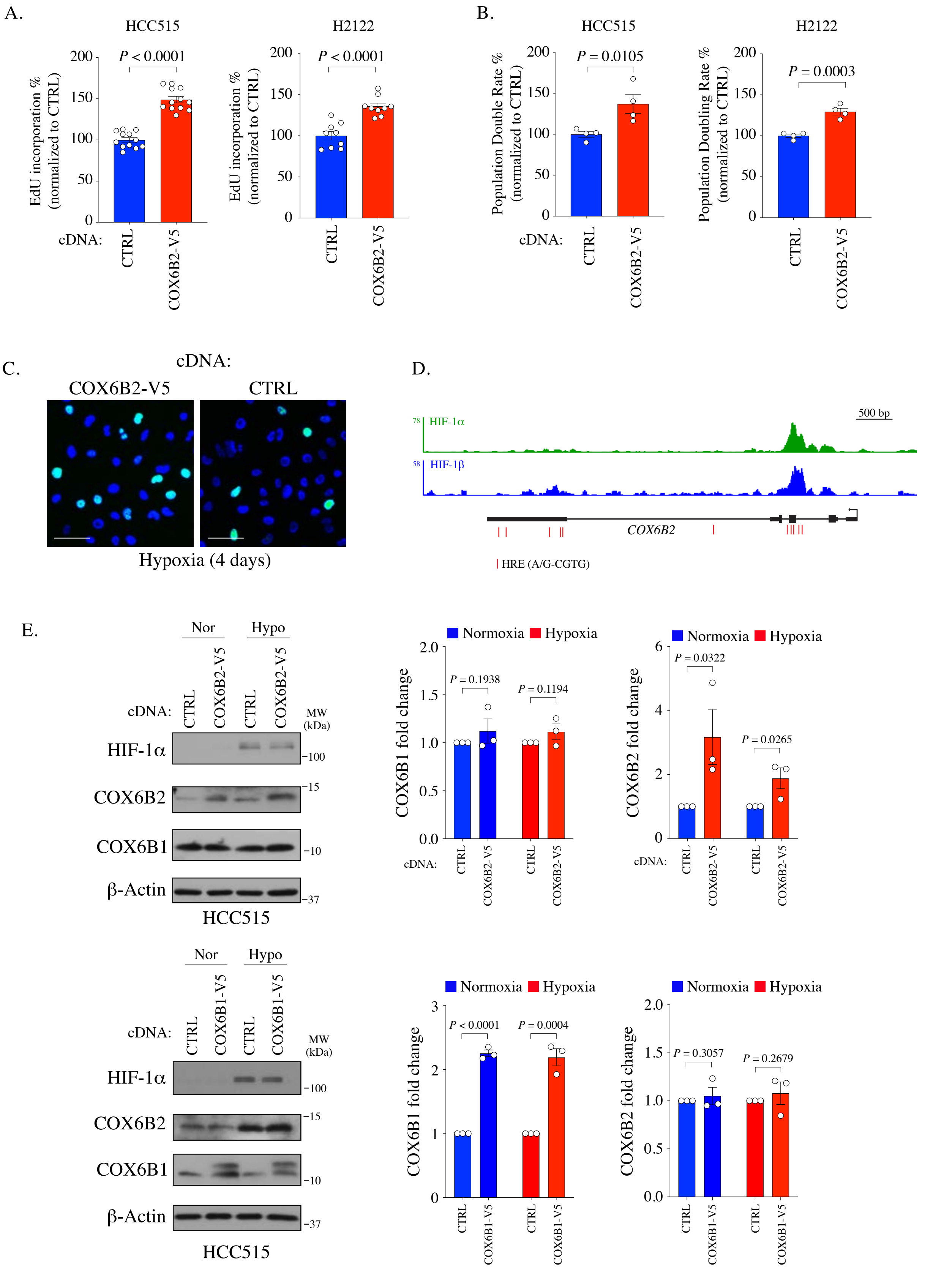

A.
HCC515
H2122
P < 0.0001
P < 0.0001
EdU incorporation % (normalized to CTRL)
cDNA:
CTRL
COX6B2-V5
B.
HCC515
H2122
P = 0.0105
P = 0.0003
Population Double Rate % (normalized to CTRL)
Population Doubling Rate % (normalized to CTRL)
C.
cDNA:
COX6B2-V5
CTRL
Hypoxia (4 days)
D.
500 bp
HIF-1α
HIF-1β
COX6B2
HRE (A/G-CGTG)
E.
Nor
Hypo
MW (kDa)
HIF-1α
COX6B2
COX6B1
β-Actin
HCC515
Normoxia
Hypoxia
P = 0.1938
P = 0.1194
COX6B1 fold change
P = 0.0322
P = 0.0265
COX6B2 fold change
COX6B1-V5
P < 0.0001
P = 0.0004
P = 0.3057
P = 0.2679

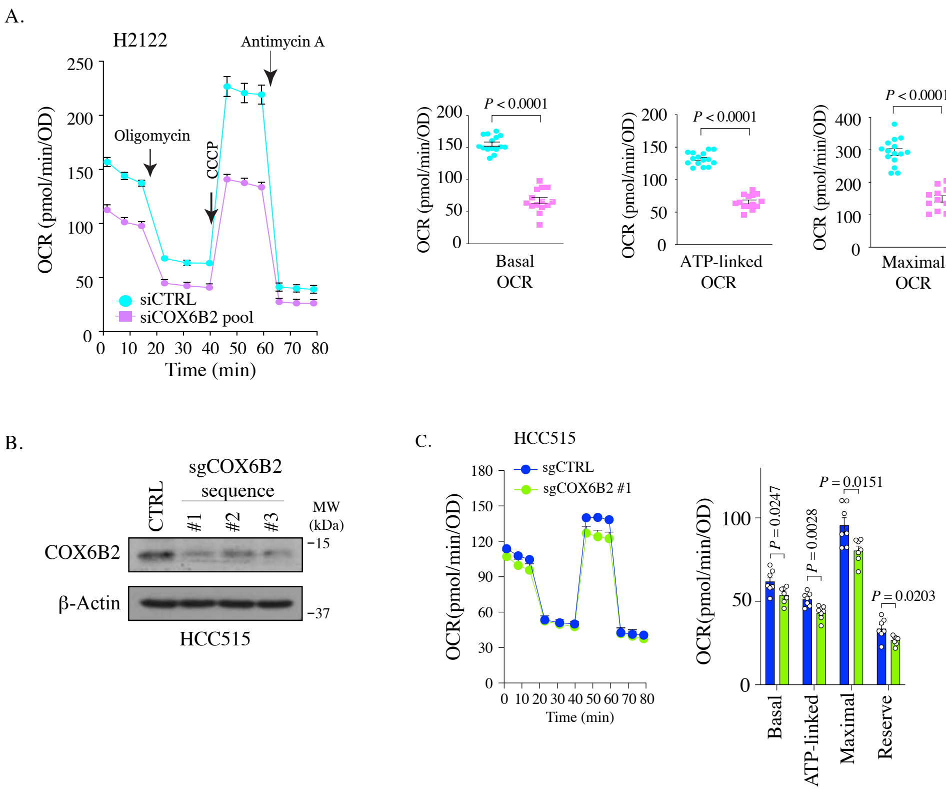
A.
H2122
Antimycin A
Oligomycin
CCCP
OCR (pmol/min/OD)
siCTRL
siCOX6B2 pool
Time (min)
P < 0.0001
Basal OCR
ATP-linked OCR
Maximal OCR
Reserve Capacity
B.
sgCOX6B2 sequence
CTRL
#1
#2
#3
MW (kDa)
COX6B2
β-Actin
15
37
HCC515
C.
HCC515
sgCTRL
sgCOX6B2 #1
OCR(pmol/min/OD)
Time (min)
P = 0.0247
P = 0.0028
P = 0.0151
P = 0.0203
Basal
ATP-linked
Maximal
Reserve

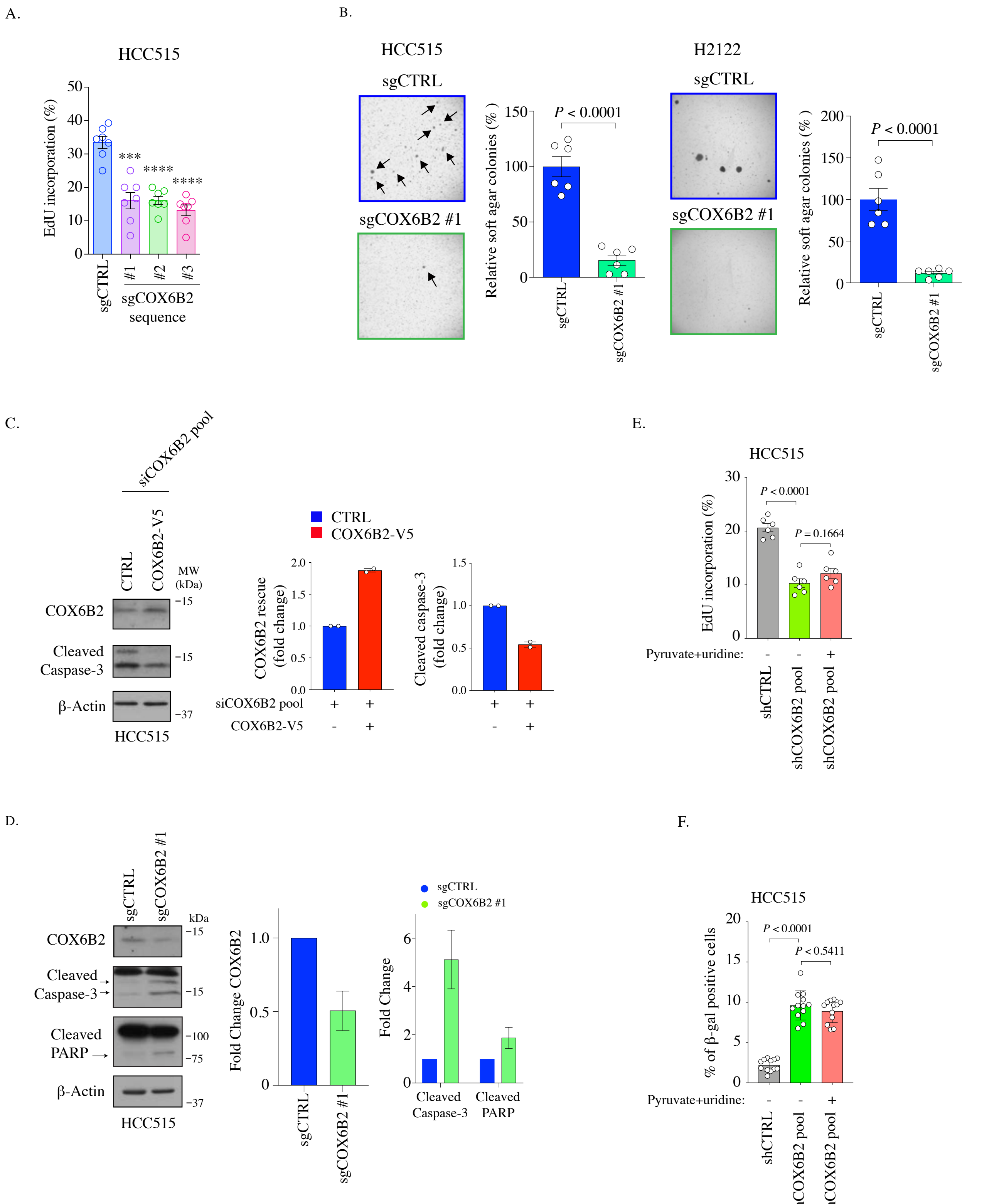
A.
HCC515
EdU incorporation (%)
sgCTRL
#1
#2
#3
sgCOX6B2 sequence
B.
HCC515
sgCTRL
sgCOX6B2 #1
Relative soft agar colonies (% )
P < 0.0001
H2122
C.
siCOX6B2 pool
CTRL
COX6B2-V5
MW (kDa)
COX6B2
Cleaved Caspase-3
β-Actin
HCC515
COX6B2 rescue (fold change)
Cleaved caspase-3 (fold change)
siCOX6B2 pool
D.
kDa
Cleaved PARP
Fold Change COX6B2
Fold Change
E.
P = 0.1664
Pyruvate+uridine:
shCTRL
shCOX6B2 pool
F.
% of β-gal positive cells
P < 0.5411

**Figure 1-figure supplement 1. COX6B2 expression in LUAD. (A)** Z-score of caspase3/7 activity screening in indicated cell lines based on data generated in (Maxfield et al. 2015). **(B)** Top: Whole cell lysates from HCC515 depleted or COX6B2-V5 or COX6B1-V5 overexpressing cells were immunoblotted with indicated antibodies. MW markers are indicated. Bottom: Graphs COX6B1 and COX6B2 protein expression from immunoblots above. Bars represent mean ± SEM (n = 3). *P* values calculated by Student's *t*-test. **(C)** Clinical data associated with tumors used in Figure 1E. **(D)** Pearson's correlation analysis of COX6B2 versus COX6B1 and COX6B2 versus COXIV protein expression in LUAD tumors as measured by immunoblots in Figure 1E. Values are relative to loading control (β-Actin).

**Figure 3-figure supplement 1. Measurement of complexes and supercomplexes. (A)** Left: Representative immunofluorescence images of COXIV in HCC515 cells +/- COX6B2-V5. Scale bar, 20 μm. Right: Quantification of staining intensity in indicated cell lines. Bars represent mean ± SEM (n ≥ 35). *P* value calculated by Student's *t*-test. **(B)** BN-PAGE gel immunoblotted with anti-COX6B2, anti-COXIV (indicating complex IV), anti-UQCRC2 (indicating complex III), and anti-NDUFA9 (indicating complex I) in HCC515 cells +/- COX6B2-V5. Representative image of n ≥ 3.

**Figure 4-figure supplement 1. COX6B2 promotes cell division. (A)** Relative EdU incorporation (%) in indicated samples. Values represent the mean ± SEM (n ≥ 9). *P* value calculated by Student's *t*-test. **(B)** Relative population doubling rate (%) in indicated samples (no pyruvate). Bars represent means ± SEM (n = 4). *P* value calculated by

Student's *t*-test. **(C)** Representative images of EdU staining in indicated cells under hypoxia for 4 days (EdU, green; Hoechst, blue). Scale bar, 100 μm. **(D)** ChIP-seq data showing HIF-1 peaks in COX6B2 in T47D cells (breast cancer). Red lines indicate HREs (Zhang et al. 2015). **(E)** Left: Whole cell lysates from indicated cell lines were immunoblotted with indicated antibodies. Hypoxia exposure was for 12 hours. Representative immunoblots from n = 3. MW markers are indicated. Nor, normoxia; Hypo, hypoxia. Right: Quantitation of protein expression. Bars represent mean ± SEM (n = 3). *P* value calculated by Student's *t*-test.

**Figure 5-figure supplement 1. Depletion of COX6B2 impairs OXPHOS. (A)** Left: Oxygen consumption rate (OCR) as a function of time in siCOX6B2 depleted H2122 cells following exposure to electron transport chain complex inhibitors. Bars represent mean ± SEM (n ≥ 14). Right: Bars represent mean ± SEM 20 minutes following the addition of each drug (on left). *P* values calculated by Student's *t*-test. **(B)** Representative images of western blots from whole cell lysates of indicated cells. **(C)** OCR was measured as a function of time for each parameter in indicated cells. Bars represent mean ± SEM (n = 7). *P* value calculated by Student's *t*-test.

**Figure 6-figure supplement 1. Phenotypes following COX6B2 depletion. (A)** The percentage of EdU-positive cells of COX6B2 depleted by three different sgRNAs in HCC515 cells. Bars represent mean ± SEM (n = 7). *P* value calculated by Student's *t*-test. ****P* < 0.001, *****P* < 0.0001. (**B)** Representative images of colonies in soft agar. Arrows indicate formed colonies. Graphs represent relative colony numbers in indicated cells. Bars represent mean ± SEM (n = 6). *P* value calculated by Student's *t*-test. **(C)** Left: whole cell lysates from HCC515 +/- COX6B2-V5 were immunoblotted with indicated antibodies. MW markers are indicated. Right: quantitation of immunoblots on left. Bars represent mean ± range (n = 2). **(D)** Representative images and quantitation of western blots from whole cell lysates of indicated cells and immunoblotted with indicated antibodies. Bars represent mean ± SD (n = 3). **(E)** The percentage of EdU-positive cells of indicated cells cultured with or without pyruvate (1 mM) and uridine (100 μg/ml) for eight days. Bars represent mean ± SEM (n = 6). *P* value calculated by Student's *t*-test. **(F)** Quantification of senescence-associated- β-galactosidase positive cells in indicated cells cultured with or without pyruvate (1 mM) and uridine (100 μg/ml) for 11 days. Bars represent mean ± SEM (n = 12). *P* calculated by Student's *t*-test.
